# 2.8 Å resolution cryo-EM structure of human parechovirus 3 in complex with Fab from a neutralizing antibody

**DOI:** 10.1101/410217

**Authors:** Aušra Domanska, Justin W. Flatt, Joonas J.J. Jukonen, James A. Geraets, Sarah J. Butcher

## Abstract

Human parechovirus 3 (HPeV3) infection is associated with sepsis in neonates characterized by significant immune activation and subsequent tissue damage. Strategies to limit infection have been unsuccessful due to inadequate molecular diagnostic tools for early detection and lack of a vaccine or specific antiviral therapy. Towards the latter, we present a 2.8 Å-resolution structure of HPeV3 in complex with fragments from a neutralizing human monoclonal antibody AT12-015 using cryo-EM and image reconstruction. Modeling revealed that the epitope extends across neighboring asymmetric units with contributions from capsid proteins VP0, VP1, and VP3. Antibody decoration was found to block binding of HPeV3 to cultured cells. Additionally at high-resolution, it was possible to model a stretch of RNA inside the virion and from this identify the key features that drive and stabilize protein-RNA association during assembly.

**Importance**

HPeV3 is receiving increasing attention as a prevalent cause of sepsis-like symptoms in neonates, which despite the severity of disease, there are no effective treatments available. Structural and molecular insights into virus neutralization are urgently needed, especially as clinical cases are on the rise. Towards this goal, we present the first structure of HPeV3 in complex with fragments from a neutralizing monoclonal antibody. At high-resolution it was possible to precisely define the epitope that when targeted, prevents virions from binding to cells. Such an atomic-level description is useful for understanding host-pathogen interaction, viral pathogenesis mechanisms, and for finding potential cures for infection and disease.

## Introduction

HPeV3 is a small, non-enveloped, single-stranded, positive-sense RNA virus, belonging to the *Parechovirus* genus of *Picornaviridae*, which currently includes 19 genotypes most commonly associated with mild gastrointestinal and respiratory illness (1, 2). Increased availability of sequence data in clinical settings has clarified that HPeV3 causes the most virulent infections of the HPeVs, particularly in infants less than 3 months of age where sickness can trigger a sepsis-like dysregulated host response often involving the central nervous system (3-9). In cases of acute meningitis or encephalitis where patients may develop abnormal white matter lesions, neurological sequelae and even death may occur (10-15). To date, no effective treatments for HPeV3 infection are available, highlighting the urgent need for a greater understanding of the structural and molecular basis for HPeV3 neutralization, especially as epidemics are likely to continue (2, 16-18). The HPeV3 virion is composed of 60 copies of the three structural proteins (VP0, VP1, and VP3) that fit together to form a 28-nm-diameter icosahedral shell around the ∼7.3 kb single-stranded RNA viral genome (19). The genome encodes a single polyprotein that during infection is subsequently cleaved into all the essential capsid components and replication proteins (2A, 2B, 2C, 3A, 3B, 3C and 3D) (20). In HPeV1, recent work has shown that newly synthesized viral RNA contains ∼60 spatially defined, conserved sequence/structure GXUXUXXU motifs that bind capsid proteins, driving genome encapsidation and efficient capsid self-assembly (21, 22). Assembled capsids lack cleavage of VP0 into VP2 and VP4 products, resulting in a shell made of three proteins rather than the four found in most other picornaviruses. Around the pentamers there is a depression referred to as the canyon. The tips of the three-fold symmetric propeller-like protrusions are adjacent to this canyon.

The VP1 C terminus of several human parechoviruses (e.g., HPeV -1, -2, -4, and -5) contains an arginine-glycine-aspartic acid (RGD) motif that can attach to αVβ1, αVβ3, and αVβ6 integrin receptors (23, 24). HPeV3 lacks the RGD motif and thus likely uses a different, as yet unknown, receptor for cell entry (25). Reliance on a different receptor may alter tissue tropism and could explain why HPeV3 infections have different clinical and epidemiological features compared to other HPeV genotypes.

Human monoclonal antibodies (mAbs) can be exploited to gain valuable insights into the structural basis for neutralizing activity, which in turn can be used for developing effective treatments. High-resolution mapping of mAb sites at the HPeV3 capsid surface allows for identification of epitopes recognized by the humoral immune system and may begin to provide mechanistic clues into immune surveillance, evasion, or escape. Here, using high-resolution cryo-EM we define such an epitope, that when targeted by a human monoclonal antibody, blocks attachment of virions to host cells; hence also describing a potential site for receptor binding.

## Results

### Cryo-EM structure of the HPeV3-Fab AT12-015 complex

Cryo-grids containing vitrified Fab-labeled virus were imaged and after data processing in RELION, a total of 74,927 particle projections were selected, which yielded a 3D reconstruction extending to 2.8 Å resolution according to the gold-standard Fourier shell correlation 0.143 criterion (Table 1 and Figure 1A) (26). Fab decoration on the capsid surface helped to assign particle orientations during data processing. The capsid was resolved to 2.3 Å, whereas small stretches of capsid associated RNA and peripheral regions of Fab were defined at a resolution lower than 3.5 Å (Figure 1B). In the structure, Fab molecules bind to symmetry-related sites at the tips of the propellers on the surface of the virion (Figure 2A-C). For fitting, large regions of the map showed clear delineation of secondary structural elements, including amino acid and nucleic acid densities (Figure 2D-G). In this manner, we could accurately map the antibody footprint on the capsid surface, as well as visualize an RNA base-stacking motif that stably anchors the genome to the capsid via interaction with a tryptophan residue (Trp 24) in the HPeV3 VP3 coat protein (Figure 2F and G).

**Table 1:** Summary of cryo-EM data collection, refinement, and validation statistics

| HPeV3-Fab complex |  |
| --- | --- |
| **Data collection** |  |
| Voltage (kV) | 300 |
| Electron exposure (e-/Å<sup>2</sup> X s) | 48 |
| Pixel size (Å) | 1.06 |
| Number of micrographs | 6,541 |
| **Reconstruction** |  |
| Number of particles | 74,927 |
| B factor (Å<sup>2</sup>) | -70 |
| FSC threshold | 0.143 |
| Resolution (Å) | 2.8 |
| **Model building** |  |
| VP0 (amino acid coverage) | 34 - 289 |
| VP1 (amino acid coverage) | 25 - 219 |
| VP3 (amino acid coverage) | 16 - 256 |
| RNA (nt) | 8 |
| Fab heavy chain (V<sub>H</sub>) | 2 - 119 |
| Fab light chain (V<sub>L</sub>) | 2 - 109 |
| **Model validation** |  |
| MolProbity score | 1.55/96<sup>th</sup> percentile<br>(protein) |
| Ramachandran outliers (%) | 0.93 (protein) |
| Poor rotamers (%) | 3.91 (protein) |
| Clashscore | 0 (protein), 4.35 (RNA) |

**Figure 1:**
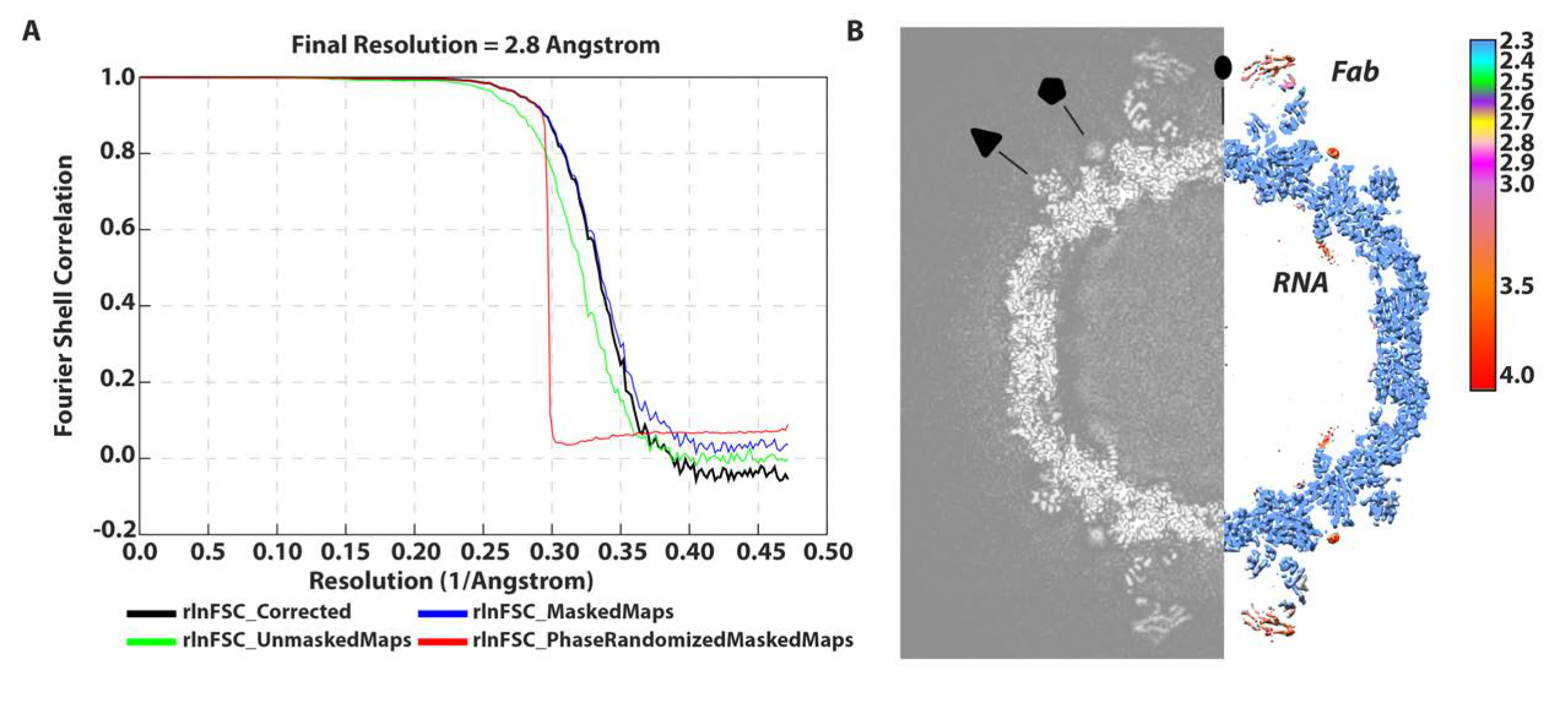
Resolution assessment for the HPeV3-Fab complex. **(A)“**Gold standard” FSC curve showing an overall nominal resolution at 2.8 Å. (B) Central cross-section of three-dimensional density map alongside structure colored according to local resolution. Two-, three- and five-fold symmetry axes are labeled.

**Figure 2:**
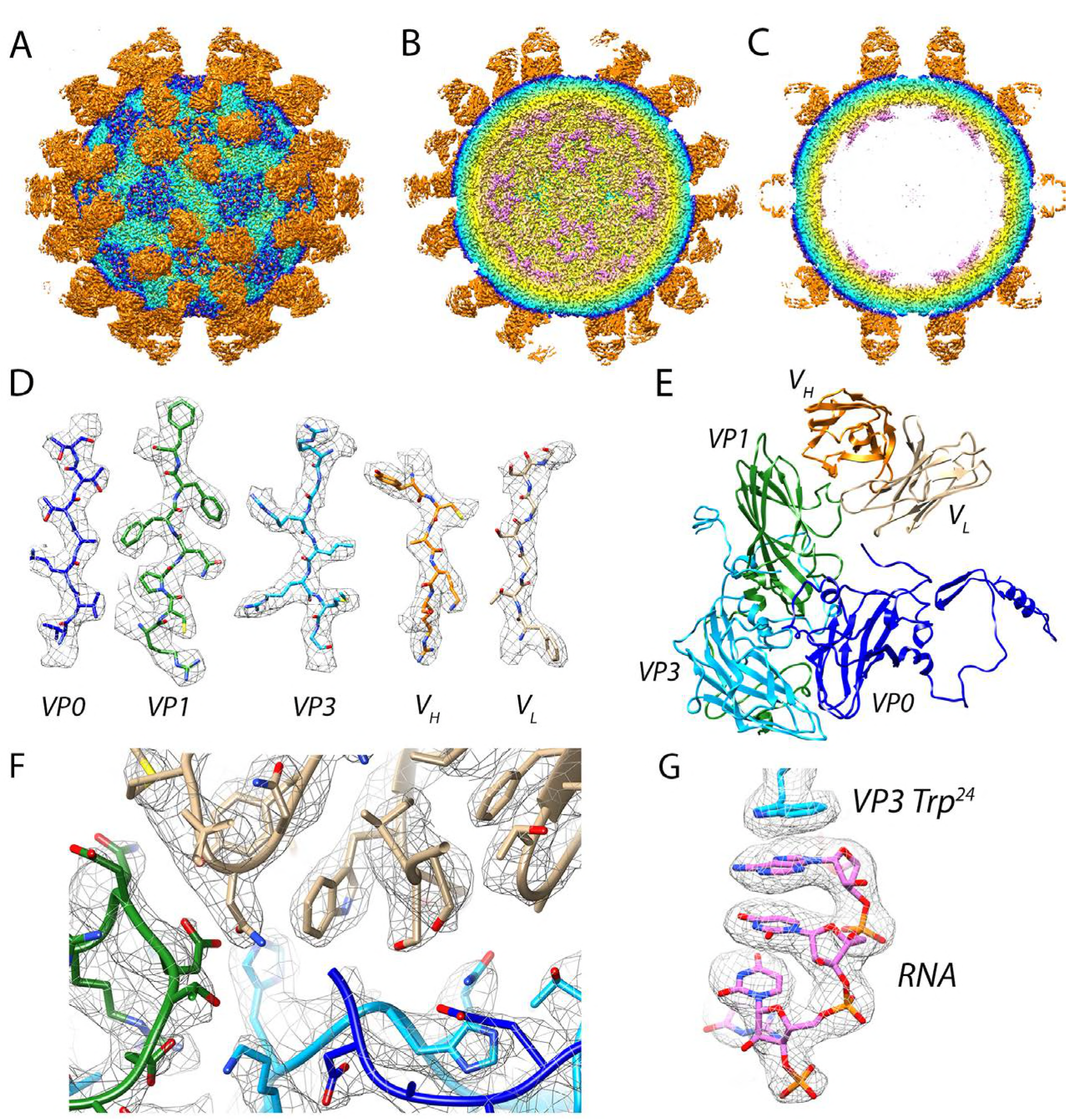
Visualization of Fab-decorated HPeV3. (A-C) Virus-Fab reconstruction shown radially color-coded (purple 100 Å, yellow 120 Å, light blue 140 Å, blue 145 Å, and orange 150 Å). (A) Surface view down a two-fold axis of symmetry. Propeller regions can be seen as a dark blue triangles on the capsid surface, the Fabs are orange. (B) Cutaway view showing RNA (purple) at five-fold vertices inside the viral capsid. (C) Central cross-section with well-defined layers of density corresponding to RNA, capsid, and Fab. (D) Side chains for viral coat proteins VP0 (blue), VP1 (green), and VP3 (light blue), as well as Fab heavy and light chains V_H_ (orange) and V_L_ (light brown) respectively. (E) Modeled asymmetric unit with a Fab molecule bound. (F) High resolution at the Fab AT12-015-HPeV3 interface. (G) RNA anchoring on the inner surface of the virus is mediated by a tryptophan (Trp 24) residue from VP3.

### Characterization of the Fab AT12-015 Epitope

In the reconstruction, the signal related to the Fab heavy and light chains is roughly as strong as that of the viral capsid, indicating 100% occupancy of the 60 available binding sites on the virion (Figure 1B). The quality of the map was such that we could fit atomic coordinates for the Fab, as well as for the three viral coat proteins, and this was followed by MDFF all-atom refinement. Results from flexible fitting revealed that the Fab targets an extended, solvent-accessible VP0-VP1-VP1´-VP3´ (´ denotes a neighboring asymmetric unit) conformational epitope. There is no evidence of induced structural changes in any of the capsid proteins upon Fab binding. Using a 3.6 Å distance cutoff we identified 28 capsid residues forming the epitope and 29 Fab residues that form the paratope (Figure 3A). Residues from the capsid that are involved in forming the immune complex are conserved among HPeV3 strains, but not for HPeV1 or other parechovirus types. Six hydrogen bonds at the interface were identified, three from the heavy chain: Arg 58 to VP1 residue Asp 87, Tyr 59 to VP1 residue Asn 138, and Arg 99 to VP3 residue Leu 252, and three from the light chain: Ser 28 to VP0 residue Glu 285, Asn 93 to VP1 residue Asp 137, and Ser 30 to VP3 residue Gly 207, that have angles in the range of 138-180° and are closely spaced to stably interact with exposed backbone nitrogen and oxygen atoms in the capsid proteins (Figure 3B). An additional hydrogen bond may form between the Fab light chain residue Ser 67 and VP3 residue Gln 209. However, the density in this area of the map was weak and because of this it was not possible to assess whether suitable geometric conditions were met for the interaction to occur other than the fact that the residue pair was in close proximity. One salt bridge between heavy chain amino acid Glu 105 and VP3 residue His 206 was inferred by the fact that centroids of the oppositely charged functional groups of the residues were within a 4 Å cutoff; and the Glu carbonyl oxygen atom was within 4 Å distance from the nitrogen atom of the His side chain (Figure 3B). This His 206 is centrally located in the footprint for the antibody and it was recently reported to be critical for binding and neutralization based on experimental selection of an antibody AT12-015 resistant HPeV3 variant (VP3 His 206 to Tyr) (27).

**Figure 3:**
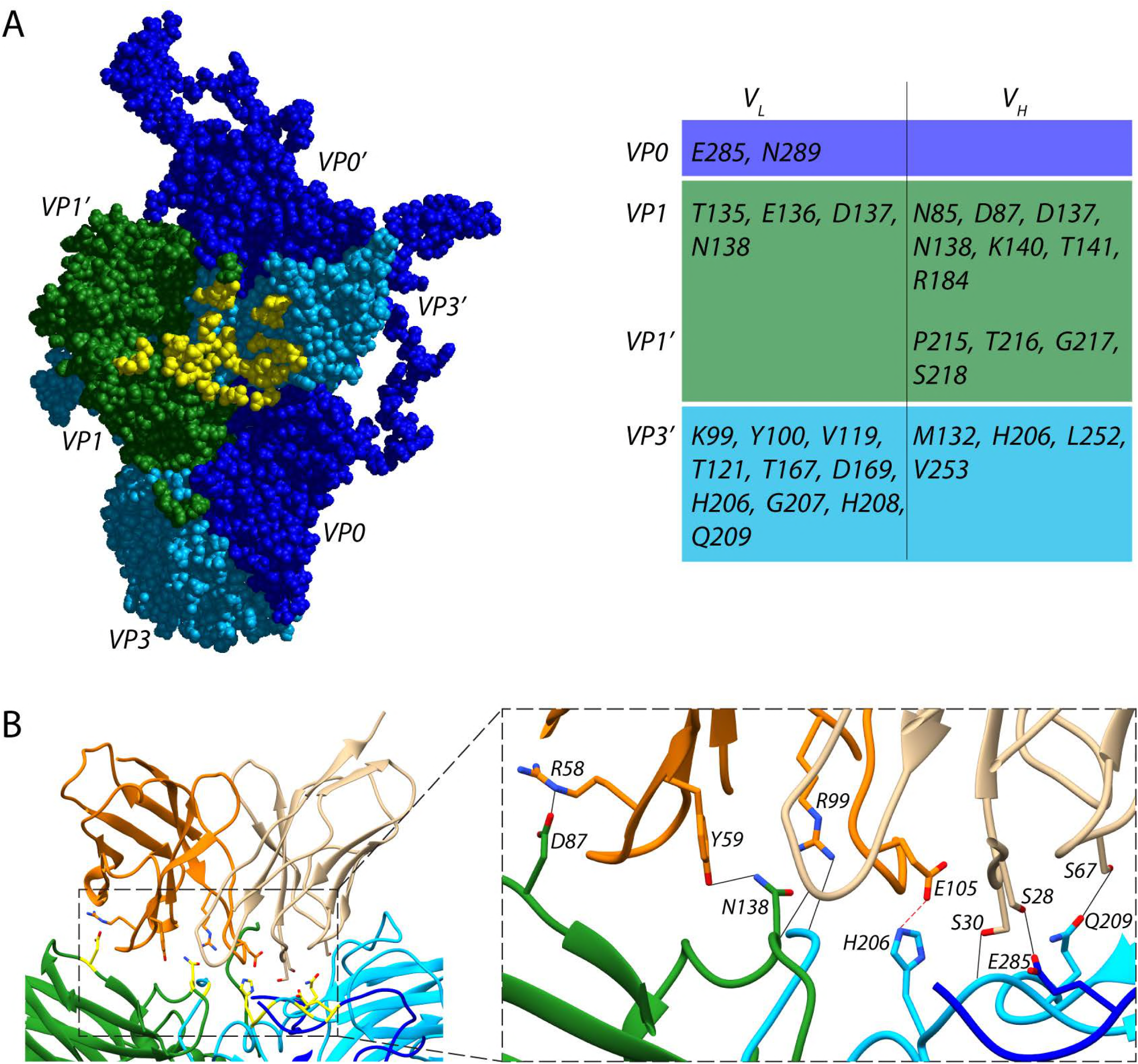
Interactions between Fab AT12-015 and HPeV3. (A) Fab binds to an epitope extended across neighboring asymmetric units in the assembled virion. Viral capsid residues that participate in Fab heavy (V_H_) and light chain (V_L_) binding are highlighted in yellow and are reported in the accompanying table. (B) Stabilizing interactions at the interface. Residues that form hydrogen bonds or a salt bridge are highlighted in yellow on the left, and colored by chain on the enlarged inset on the right. In the in-set, hydrogen bonds are shown as black dashed lines along with a centrally located salt-bridge high-lighted in red.

### Human monoclonal antibody AT12-015 prevents HPeV3 entry into cells

To probe HPeV3 A308/99-specific neutralization, we assayed whether binding of virions to human intestinal HT-29 cells was blocked by the presence of human monoclonal antibody AT12-015 (Figure 4A). For antibody-mediated blocking, we pre-incubated equivalent amounts of virus with antibody at 1:10, 1:100, 1:1000, and 1:10000 dilutions for 1 hour at 37° C and then added the complexes to cells. Cellular attachment proceeded under ice-cold conditions for 1 hour and afterwards unbound virions were removed with a series of gentle PBS wash steps. Cold binding ensures that HPeV3 remains at the surface of cells and is not internalized. When preformed antibody-decorated virions were added, either from mixing HPeV3 with stock or 1:10 dilution of antibody, no fluorescence signal was observed on the surface of HT29 cells similar to the mock infection (no virus) and antibody controls (Figure 4B - stock, 1:10, mock, antibody no virus). Small-sized clusters of virions were infrequently observed at excess levels of antibody (Figure 4B - stock, 1:10). In contrast, staining was clearly visible on the surface of cells incubated with HPeV3 alone, as well as with higher dilutions of antibody, and fluorescence intensity was weaker at 1:100 dilution albeit with a high standard deviation (Figure 4B - 1:10000, 1:1000, and 1:100). Virus mixed with 1:100 of antibody showed neutralizing activity without any apparent clumping (Figure 4B - 1:100). Taken together, these results indicate that AT12-015 neutralizes HPeV3 infection extracellularly rather than by a post-attachment mechanism.

**Figure 4:**
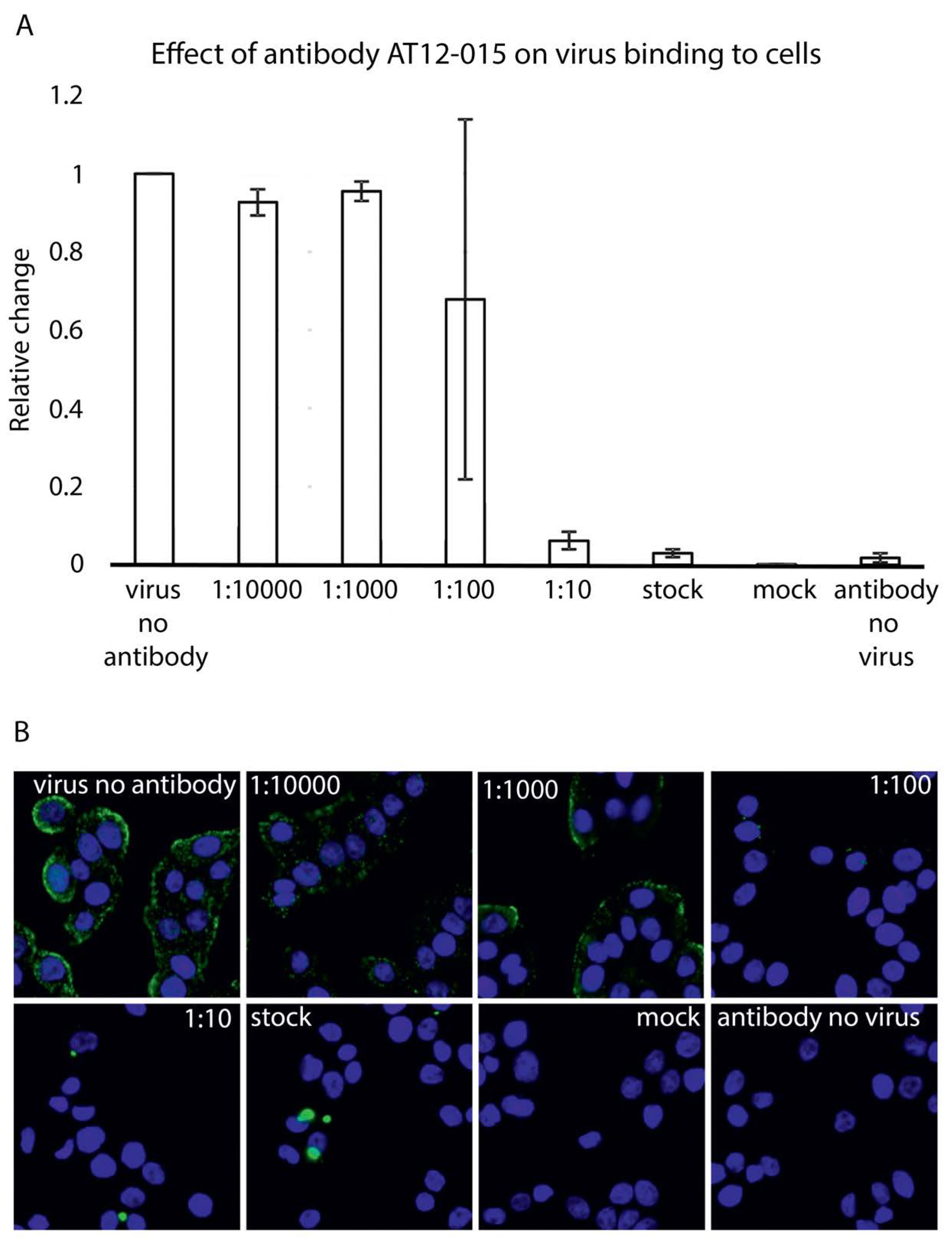
Antibody AT12-015 blocks virus binding to HT29 cells. (A) Effect of preincubation of HPeV3 with different amounts of human monoclonal antibody AT12-015. Virus was mixed with antibody for 60 minutes at 37°C and then added to HT29 cells for 60 minutes cold binding. Cell nuclei were visualized using a Hoechst stain and virus binding was scored by measuring Alexa Fluor 488 intensity. The results are the average of three repeats of the cold binding assay. The error bars represent the standard error of the mean (SEM). (B) Representative fluorescence images of HPeV3 incubated in the presence or absence of varying amounts of antibody.

### RNA inside the HPeV3 capsid

The HPeV3 virion contains ∼7.4 kb of mostly unstructured single-stranded RNA genome. In the assembled particle, roughly 25% of the RNA adopts a defined conformation near the inner capsid surface, lining symmetry-related sites directly beneath the icosahedral five-fold vertices (19). For parechoviruses, detailed structural analysis of genome-capsid interactions has only been carried out for HPeV1 (21). Here we performed a similar analysis for HPeV3. The inside of our 2.8 Å HPeV3 map shows stretches of RNA in the same region as defined for HPeV1. The RNA has a defined tertiary structure, forming a single-stranded loop with a significant portion involved in a rigid base-stacking motif that is capped by an aromatic side chain residue Trp 24 of VP3 with a small conjugated π system (Figure 5A and B). In this way, the tryptophan has stabilized its π orbitals to a resonance level with the aromatic orbitals of adjacent RNA bases to enable efficient packaging and formation of stable virions. EM density for the planar stacking profile is well-resolved, on the order of 25 Å in length, before reaching a helix-coil transition (Figure 5B). Coordinates for the six RNA nucleotides from previous structural work on the HPeV1 virion are in good agreement with our modeled stacking motif. We fitted eight bases of RNA beneath the capsid, which included a portion of the packaging sequence described recently for HPeV1 (22). The final sequence docked was U^0^-G^1^-G^2^-U^3^-A^4^-U^5^-U^6^ U^n^. Using the RNA motif we searched the HPeV3 A308/99 genome (Genbank code AB084913) and identified 33 sequences that contained G^1^-purine^2^-U^3^-purine^4^-U^5^, 13 of which include the full motif X^0^-G^1^-purine^2^-U^3^-purine^4^-U^5^-X^6^-X^7^-U^8^.

**Figure 5:**
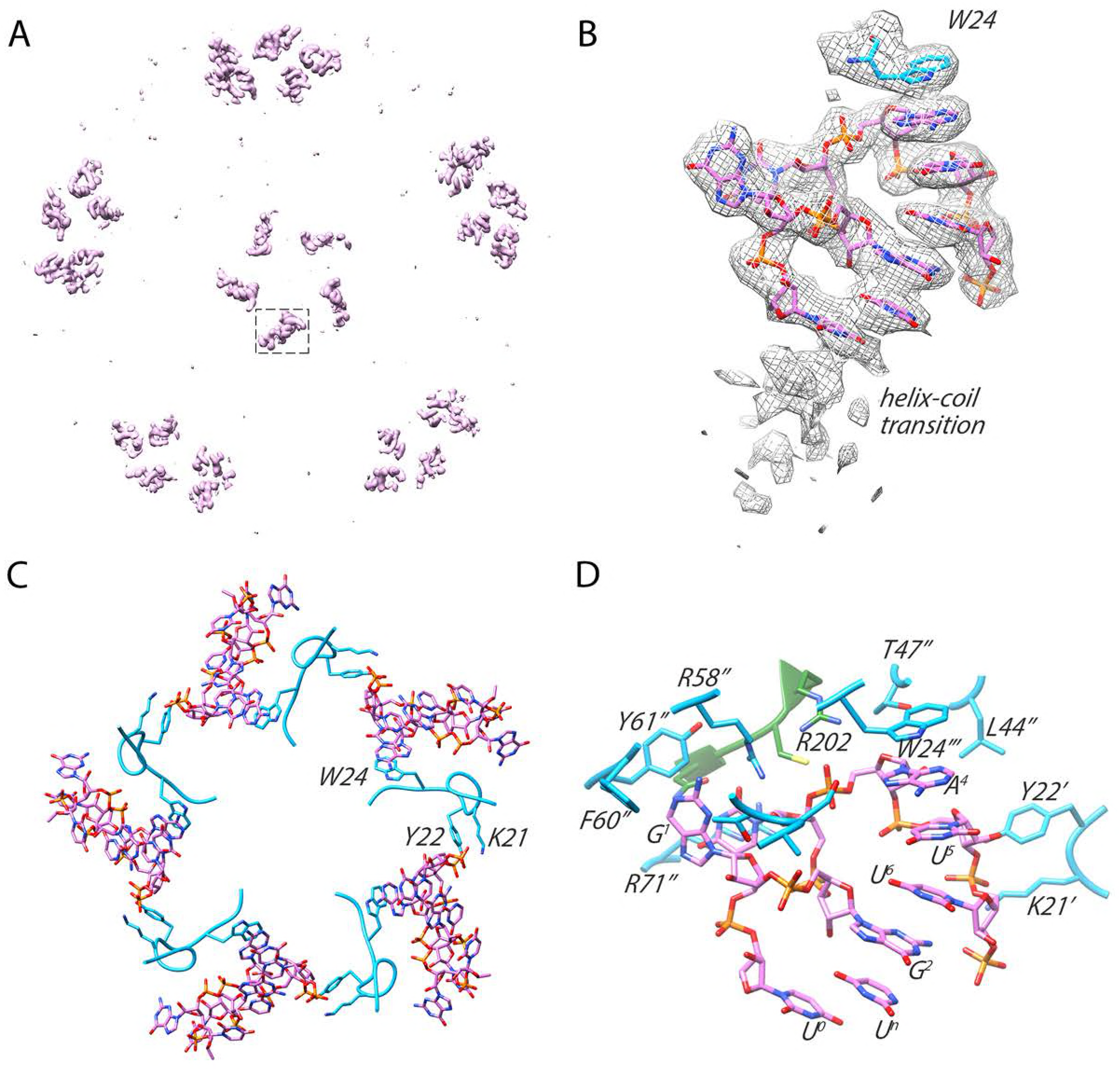
Ordered RNA inside the HPeV3 virion. (A) RNA density segmented from within the virion seen along an icosahedral five-fold axis. The boxed segment of the density is enlarged in B. (B) Eight nucleotides fit to their corresponding density before reaching the helix-coil transition. (C) VP3 tails bridge two adjacent loops of RNA to promote efficient packaging and assembly. Small portion of the capsid VP3 sequence (AAs Leu 16-Arg 26) is shown to clarify the VP3-RNA network on the inner surface of the viral capsid. (D) The binding pocket for RNA on the inside of the capsid involves residues from VP1 and VP3. One RNA loop is stabilized by residues from a single VP1 chain (green) and three VP3 chains (light blue designated by ´,´´, and ´´´).

In addition to modeling the Trp 24-RNA contact in the cryo-EM structure, two other residues from the same VP3 strand, Lys 21 and Tyr 22, were found to interact with a neighboring RNA loop. These residues, along with Trp 24, in the context of the assembled pentamer, appear to be key for stabilizing the stacked RNA below the capsid vertex (Figure 5C). In fact, mutation of two of the amino acids, either Tyr 22 or Trp 24 to alanine, in HPeV1 is lethal confirming their essential role in virion assembly (22). Other capsid residues that further support the RNA loop under the vertex were identified in VP1, as well as VP3. Specifically, VP1 residues Arg 202, Cys203, and Asn 205, and VP3 residues Ala 18, Ser 19, Thr20, Leu 44, Thr 47, Arg 58, Phe 60, Tyr 61, and Arg 71, many of which are aromatic or positively charged, form important RNA base and backbone contacts, as well as complement the negative charge of the RNA (Figure 5D). In HPeV1, mutations of VP1 residues Arg 202 and Cys 203 to Ala as well as VP3 residues Thr 44, Arg 55, and Arg 68 to Ala (VP3 residues Thr47, Arg 58, and Arg 71 in our structure) were shown to be lethal, indicating that these residues are important for RNA packaging and virion assembly (22).

## Discussion

Exact knowledge of the structural, antigenic, and immunogenic features of HPeV3 is essential for understanding host-pathogen interaction, viral pathogenesis mechanisms, and for finding potential cures for infection and disease, which is pressing as HPeV3 outbreaks are widespread and may cause severe sepsis-like syndrome in neonates. Recently, we determined a structure of HPeV3 in complex with Fab fragments from a human monoclonal antibody AT12-015 by cryo-EM (19). However, the strain of virus used in that study, HPeV3 isolate 152037, was not neutralized by addition of AT12-015. Furthermore the resolution of the reconstruction was only at 15 Å, which prevented atomic characterization of the epitope on the surface of the viral capsid. Here we report a 2.8 Å-resolution cryo-EM structure of an AT12-015 Fab-decorated HPeV3 virion, isolate A308/99 that is neutralized by the monoclonal antibody from infecting human intestinal HT29 cells. In the high-resolution structure, the antigen-antibody interface was well-defined and modeling of viral coat proteins and Fab molecules into density revealed an extended conformational epitope across the interface of adjacent asymmetric units, involving residues from different parts of neighboring VP0, VP1, and VP3 chains spatially juxtaposed by the structure of the capsid. When bound, monoclonal AT12-015 prevented virus attachment to target HT29 cells except for at high dilutions of antibody, indicating that neutralization occurs extracellularly and not by a post-attachment mechanism. In this way, we were able to identify the epitope of AT12-015 and determine how the antibody works against HPeV3 A308/99. In addition, the 2.8 Å map provided the most complete picture yet of ordered RNA on the inside of a human parechovirus.

Antibody AT12-015 was first isolated from the immune repertoire of a person with HPeV3 infection using the AIMSelect method and it broadly recognizes strains of HPeV3 but only neutralizes the prototype A308/99 virus that we used in the study (19, 27, 28). At high resolution we could clearly specify the conserved conformational epitope shared among HPeV3 strains, including solvent-accessible atoms in the interface region, which now includes contributions from VP0 (Glu 285, Asn 289), VP1 (Asn 85, Asp 87, Thr 135-Asn 138, Lys 140, Thr 141, Arg 184), VP1’ (Pro 215-Ser 218), and VP3’ (Lys 99, Tyr 100, Val 119, Thr 121, Met 132, Thr 167, Asp 169, His 206-Gln 209, Leu 252, Val 253; Figure 3A). No structural changes were induced upon complex formation. Atomic-level characterization brings clarity to exactly how Fab AT12-015 binds to the virion. Of particular interest is His 206 of VP3 as it was recently shown that an HPeV3 A308/99 variant mutated at this residue position to Tyr escapes neutralization by AT12-015. Our results provide important context to this observation by showing that the nitrogen atom of VP3 His 206 forms a salt bridge with the carbonyl oxygen of Glu 105 in the Fab heavy chain at the center of the antibody footprint (27). Thus we can confirm a key role for VP3 His 206 in driving complex formation.

Because antibodies often prevent virus attachment and entry into target cells, we tested whether AT12-015 blocks HPeV3 A308/99 binding to human intestinal HT29 cells. We found that mixing antibody with virions under saturating conditions efficiently inhibits viral adhesion to cells and this effect is only reversed at high dilutions of the antibody. In the presence of high amounts of AT12-015 small clumps of virus particles were infrequently observed, indicating that antibody-mediated aggregation plays a minor role in exacerbating adherence. Neutralization occurs thus presumably by either directly or indirectly preventing receptor engagement. Such specific targeting rather than generalized clumping is supported by the fact that of all HPeV3 strains bound by AT12-015, only isolate A308/99 is neutralized (27). To date, little is known about HPeV3 receptor and co-receptor dependencies as it lacks the RGD motif and hence probably utilizes an uptake mechanism that differs from other parechovirus types. This could account for type-specific neutralization, and in general usage of a different receptor by HPeV3 likely influences tropism and thus the unique disease severity in the human population.

It has become increasingly appreciated in recent years that for single-stranded RNA viruses, the life cycle is in part regulated by the secondary and tertiary structure of their genomes, hence the current high priority to understand protein-(single-stranded) RNA recognition motifs and RNA sequence-specific folding as it occurs in assembled virions (22, 29-33). Such information may help with efforts to inhibit viral propagation, as well as in nanotechnology applications aimed at harnessing either virus or virus-like systems for efficient gene delivery (34). Based on our cryo-EM data, we propose a mechanism for adding HPeV3 RNA to assembling shells that utilizes π electron delocalization from VP3 side chain Trp 24 to assist with nucleotide binding. Here, at the mechanistic level, RNA folding is stabilized by a capsid-locking step where Trp 24, accessible on the inner surface of capsid protein VP3, forms a geometrically favorable short-range stacking interaction with a purine, which is then further strengthened by long-range interactions as a result of electronic resonance through further stacking of adjacent nucleotides. The 14 additional highly conserved capsid residues: VP1 Arg 202, Cys 203 and Asn 205, VP3 Ala 18, Ser 19, Thr 20, Lys 21, Tyr 22, Leu 44, Thr 47, Arg 58, Phe 60, Tyr 61, and Arg 71 help to position the ordered RNA loop against the inside of the capsid. From an evolutionary perspective, this means that over time the inner surface of the virus has been fine-tuned to orchestrate the interaction between VP3 Trp 24 and discrete sequences of genomic RNA, and that likewise the spacing of recognition motifs on the HPeV3 genome has been thermodynamically optimized so as to minimize the free energy of capsid assembly.

## Methods

### Virus sample preparation

Human colon adenocarcinoma (HT29) cells were propagated in McCoy’s 5A medium supplemented with 1X non-essential amino acids, 1X antibiotic-antimycotic, and 10% fetal bovine serum with the culture condition of 37°C and 5% CO_2_. Cells were grown to ∼90% confluency before inoculating with human parechovirus 3 (HPeV3) isolate A308/99 at a multiplicity of infection of 0.1. HPeV3 A308/99 was grown in fresh medium as described above except that the medium was slightly modified to contain 1 mM MgCl_2_, 20 mM HEPES pH 7.4 and no FBS. Inoculated cells were incubated at 37°C for 3 days. Cells and medium were harvested. At this point concentration of HEPES was increased to 40 mM final concentration. The cells were opened by three freeze-thaw cycles and virus-containing medium was clarified by low-speed centrifugation. Afterwards the supernatant was carefully removed and concentrated via ultrafiltration using Centricon units with a cut-off at 100 kDa in weight. For purification, we applied a CsCl density gradient (top - 1.2502 g cm^-3^, bottom - 1.481 g cm ^-3^) combined with ultracentrifugation (32,000 rpm, 4°C) for 18 hours in a Beckman type SW41 Ti rotor. The virus band was collected and the buffer exchanged into 1X TNM buffer: 10 mM Tris-HCl, pH 7.5, 150 mM NaCl, 1 mM MgCl^2^. Ultracentrifugation and buffer exchange were repeated. Concentration was estimated by Coomassie-blue-stained SDS-PAGE gel, where different concentrations of bovine serum albumin solution were used as standards. Infectivity was measured using a TCID50 endpoint dilution assay.

### Generation of antigen-binding Fab fragment

HPeV3 A308/99-specific monoclonal antibody (AT12-015) was obtained from AIMM Therapeutics (the Netherlands). AT12-015 antibody was digested to produce antigen-binding Fab fragments using the Pierce Fab micro preparation kit (Pierce). The Fab concentration was assessed by Coomassie-blue-stained SDS-PAGE gel, where different concentrations of bovine serum albumin solution were used as standards. For complex formation, 30 µL of 0.1 µg/µL virus and 9 µL of 0.1 µg/µL antibody were mixed (1:60 molar ratio) and incubated for 1 hour at 37°C.

### Cryo-EM data acquisition

Sample volumes of 3 µL of purified HPeV3 A308/99-Fab complex were applied to glow-discharged ultrathin carbon-coated lacey 400-mesh copper grids (Ted Pella product #01824) and vitrified using a custom-made manual plunger. Cryo-grids were visualized with a FEI Titan Krios electron microscope operating at 300 kV accelerating voltage, at a nominal magnification of 75,000× using a FEI Falcon II direct electron detector, corresponding to a pixel size of 1.06 Å on the specimen level. In total, 6,541 images with defocus values in the range of -0.5 to -2.5 µm were recorded in movie mode with 1 second of total acquisition time. Each movie contained 18 frames with an accumulated dose of about 48 electrons per Å^2^.

### Image processing and 3D reconstruction

Dose-fractionated image stacks containing frames from 2 to 17 were subjected to beam-induced motion correction using MotionCor2 (35). Estimation of contrast transfer function parameters for each micrograph was done using Gctf (36). Particle selection, 2D classification, and 3D classification were performed on an unbinned dataset (1.06 Å/pix, 480 pixel box size) using RELION 2.0 (26). In total, 217,212 particle projections were selected. After reference-free 2D classification in RELION in the best classes containing 179,457 particle projections were used for further processing. A ∼ 10 Å reference map generated in AUTO3DEM from a modest-sized dataset of 2050 particle images collected on a FEI Tecnai TF20 cryo-electron microscope filtered to 60 Å was used for initial maximum-likelihood-based 3D classification (37). Three classes accounting for 74,927 particles were selected for 3D refinement and reconstruction. During post-processing step in RELION the map was masked with a soft mask and sharpened using a B-factor -70 Å^2^. The final refinement resulted in a 2.8 Å map based on the gold-standard Fourier shell correlation 0.143 criterion. Local resolution was determined using ResMap with the unsharpened map as an input (38).

### Atomic model building and refinement

An initial atomic model for the HPeV3 A308/99-Fab complex was generated using I-TASSER and SWISS-MODEL based on the crystal structure of the HPeV1 virion (PDB ID: 4Z92) and Fab fragments of human monoclonal antibody AM28 (PDB ID: 4UDF) (21, 39-41). Docking of atomic coordinates was done manually using UCSF Chimera and the fit was further optimized using the ‘Fit in Map’ command (42). Inspection and further refinement was done using Coot 0.8.8 and this served as input for molecular dynamics flexible fitting (MDFF) (43). The MDFF program was used together with NAMD and VMD to further enhance the fit of models into cryo-EM density (44-46). A scale factor of 1 was employed to weigh the contribution of the cryo-EM map to the overall potential function used in MDFF. Simulations included 20,000 steps of minimization and 100,000 steps of molecular dynamics under implicit solvent conditions with secondary structure restraints in place. To achieve the best fit of the model to the cryo-EM density three iterations between Coot and MDFF were performed with the last step being relaxation of the structure by an energy minimization step using MDFF. For hydrogen bond detection at the virus-antibody interface we examined structure in UCSF Chimera using a strict distance cutoff of 3.6 Å and for geometrical constrains we only included hydrogen bonds within the range of 138 - 180° (47, 48). RNA-protein interface was analyzed in UCSF Chimera using the same (3.6 Å) distance cutoff.

### Binding assay

HT29 cells were seeded on 96-well plates at a density of 40000 cells per well in the same culture conditions as during virus sample preparation. Antibody AT12-015 was incubated as stock (0.5 mg/ml) or as a dilution (1:10, 1:100, 1:1000, 1:10000) with 1x CsCl-gradient purified HPeV3. Specifically, 2 µL of antibody was mixed with 2 µL of virus (8 * 10^5 pfu/ml) for 1 hour at 37 °C. The incubation took place in McCoy’s 5A medium supplemented with 1X GlutaMAX, 1X non-essential amino acids, 1X antibiotic-antimycotic, 20 mM HEPES, and 30 mM MgCl^2^. Plates containing cells and tubes containing antibody-virus complexes were placed on ice and allowed to cool. Growth medium of the cells was exchanged to cold-binding medium. Antibody-virus complexes, as well as either viruses or antibodies alone were then added to cells and incubated for 1 hour at ice-cold temperature. After incubation, cells were washed 3 times with 0.5 % BSA-PBS. For wells containing antibody-decorated virus, the primary staining step was skipped. Otherwise, AT12-015 was used as a primary antibody (diluted in 0.5% BSA-PBS) for unlabeled virus or in wells designated to serve as an antibody control. After three additional washing steps, secondary antibody was added for 1 hour. A further series of washes was carried out and Hoechst (1 ug/ml) was added for visualization of cell nuclei. Wells were then washed a last time and plates were sealed.

### High content imaging and analyses

All experiments were performed in 96-well plates (Perkin Elmer) and images were acquired using the automated fluorescence microscope CellInsight from Thermo Scientific. Image analysis was completed using CellProfiler (http://cellprofiler.org)).

### Accession numbers

The final density map has been deposited to Electron Microscopy Databank (EMDB) with accession code EMD-0069 (https://www.ebi.ac.uk/ebisearch/search.ebi?db=allebi&query=emd-0069&requestFrom=searchBox). The atomic model has been deposited to Protein Databank (PDB) with accession code 6GV4 (http://www.ebi.ac.uk/pdbe/entry/pdb/6gv4).

## Acknowledgements

We thank Sergey Guryanov and Shabih Shakeel for helpful discussions. We thank Benita Löflund, Pasi Laurinmäki, Lauri Pulkkinen (University of Helsinki), and Jiri Novacek (Masaryk University), as well as Instruct-FI, the Biocenter Finland National cryo-electron microscopy and light microscopy units, Institute of Biotechnology, and the CSC-IT Center for Science Ltd. for providing technical assistance and facilities to carry out the work. We thank Hiroyuki Shimizu (National Institute of Infectious Diseases) and Katja C. Wolthers (Amsterdam Medical Center) for kindly providing HPeV3 A308/99. We thank Tim Beaumont (AIMM Therapeutics) for kindly providing the AT12-015 antibody. This work was supported by iNEXT (project number 653706), a Horizon 2020 program of the European Union (iNEXT PID:2141); CIISB research infrastructure project LM2015043 funded by MEYS CR (CF Cryo-electron Microscopy and Tomography CEITEC Masaryk University, Czech Republic); the Academy of Finland (275199 to S.J.B.), the Sigrid Juselius Foundation (S.J.B.), the People Programme (Marie Curie Actions) of the European Union’s Seventh Framework Programme (FP7/2007-2013) under REA grant agreement (PIEF-GA-2013-628150 to A.D.) and the Seventh Framework Programme of the European Union AIPP under contract PIAPP-GA-2013-612308 to S.J.B.

## Author Contributions

Conceptualization (A.D., S.J.B.); formal analysis (A.D.); data curation (A.D., S.J.B.); investigation (A.D., J.W.F., J.J.J.J.); methodology (A.D., J.W.F., J.J.J.J., J.A.G., S.J.B.); software (J.A.G.); validation (A.D., J.W.F.); visualization (A.D.); manuscript writing (J.W.F.); manuscript revision (A.D., J.A.G., S.J.B.); funding acquisition (A.D., S.J.B.); supervision (A.D., S.J.B.); project administration (S.J.B.).

